# Taxon-specific differences in C and N cycling and metabolic activity of intertidal organisms: Part A – Short-term processes

**DOI:** 10.64898/2026.02.24.707700

**Authors:** Tanja Stratmann, Dick van Oevelen, Marcel T. J. van der Meer

## Abstract

European tidal flats that host non-native *Magallana gigas* reefs contribute to several ecosystem functions. Among others, they provide a habitat for a large variety of associated fauna. However, we often lack detailed information about any trophic interactions of the associated macrozoobenthos species with the oysters, and about their role in the carbon and nutrient cycle. Therefore, we performed *ex-situ* pulse-chase tracer experiments in the Eastern Scheldt (Southwest Dutch Delta, Netherlands) in summer and autumn 2020, where we fed *M. gigas* and their associated fauna ^13^C- and ^15^N-enriched bacterioplankton while the macrozoobenthos was incubated in water containing deuterium oxide (^2^H_2_O; enrichment: 1 – 2.5%). The aim was (1) to assess differences in short-term (<12h) processing of bacterioplankton in summer and autumn, and (2) to study differences in ^2^H incorporation – a proxy for metabolic activity – of *M. gigas* and its associated fauna in summer and autumn. In summer, all macrozoobenthos species combined consumed significantly less bacterioplankton-derived ^13^C and ^15^N than in autumn, while all macrozoobenthos species combined incorporated comparable amounts of ^2^H into their tissue in both seasons. Most bacterioplankton-derived ^13^C and ^15^N was taken up by sponges (*Halichondria panicea, Hymeniacidon perlevis*), crabs (*Carcinus maenas, Eriocheir sinensis, Rhithropanopeus harrisii*), and limpets (*Crepidula fornicata*). Most ^2^H was taken up by crabs (*C. maenas, E. sinensis*), sponges (*H. perlevis*), and snails (*Littorina littorea*), implying that these species were the most metabolically active ones. Overall, the metabolic activity was linked to feeding activity in summer 2020, whereas in autumn 2020, the link was weaker and the most metabolically active species were not necessarily the species that had incorporated most ^13^C and/or ^15^N.

## Introduction

Oyster reefs provide several ecosystem functions, such as biodiversity (Guy et al., 2018; Martin et al., 2025), habitat (De Santiago et al., 2019; Chan et al., 2022), carbon and nutrient cycling (Dame et al., 1989; Kellogg et al., 2013; Westbrook et al., 2019), sediment trapping (Reidenbach et al., 2013), or wave attenuation (Morris et al., 2021; Salatin et al., 2022). Therefore, non-native Pacific oyster *Magallana gigas* reefs may partially fill an important ecological niche in European coastal waters after most reefs of the native European oyster *Ostrea edulis* disappeared over the last ∼200 years (zu Ermgassen et al., 2025). Historically, European *O. edulis* reefs provided habitat for more than 190 associated macrobenthos species that belong to 13 phyla and seven different trophic guilds/feeding types, like primary producers, deposit feeders, filter-/suspension feeders, parasites, herbivores, omnivores, and carnivores (Thurstan et al., 2024). In comparison, European *M. gigas* reefs host at least 86 associated macrobenthos species belonging to 11 phyla and eight feeding types, such as primary producers, bacterivores, deposit feeders, filter-/suspension feeders, grazer, herbivores, omnivores, and predators/scavengers (Diederich, 2005; Decottignies et al., 2007a; Kochmann et al., 2008; Markert et al., 2010; Green and Crowe, 2013; Guy et al., 2018).

For most of the associated fauna of *M. gigas*’, we miss information about their role in the oyster reef’s nutrient and carbon cycle. We mostly do not know, either, whether this associated fauna has trophic interactions with *M. gigas* or whether the fauna only uses the oyster shells as hard substrate. One reason for this lack of knowledge is the scientific approach applied: Traditionally, *M. gigas* specimens and their associated fauna were mostly studied separately, and not incubated together in their naturally occurring ‘oyster-associated fauna clusters’. For instance, (Decottignies et al., 2007a) investigated the trophic interactions between the two filter/suspension feeders *Crepidula fornicata* and *M. gigas* by collecting these specimens individually, brushing off all epibionts from the shells, and studying their natural abundance stable isotopic composition. This approach is very suitable as long as the study aims to investigate the role of individual species and not of the complex food web consisting of oysters and associated fauna. In contrast, in this study, we assessed which members of an ‘oyster-associated fauna cluster’ process bacterioplankton and are metabolically active, which we defined as “chemical changes that occur in a living animal […] as a result of metabolism” (Park, 2012). We were able to quantify the ^13^C and ^15^N uptake and the metabolic activity (i.e., uptake of ^2^H) of the complete macrozoobenthos assemblage at near *in-situ* conditions without disturbing present trophic and non-trophic interactions via the removal of epibionts from their substrate. For this, we performed *ex-situ* triple (^13^C, ^15^N, ^2^H) pulse-chase tracer incubation experiments with *M. gigas*, the sponge *Halichondria (Halichondria) panicea*, which is a common epibiont of *M. gigas* in the Eastern Scheldt (Stratmann et al., 2025b), and further associated fauna.

The Eastern Scheldt is a semi-enclosed, 350 km^2^ large tidal bay with a tidal range of 2.9 to 3.5 m (Jiang et al., 2019) and clear seasonal trends in temperature, oxygen, and suspended particulate matter concentrations (Horn et al., 2023). The Eastern Scheldt harbors extensive shellfish cultures and fisheries concentrating on mussels (*Mytilus edulis*), cockles (*Cerastoderma edule*), and *M. gigas* (Smaal et al., 1986, 2009, 2013; Smaal and Lucas, 2000). Since *M. gigas*’ introduction to the Eastern Scheldt in 1964 (Smaal et al., 2009; Troost, 2010), its standing stock has increased steadily (Smaal et al., 2013). In the eastern part of the Eastern Scheldt, where the specimens for this study were collected, 365 ha (equivalent to 9.12% of the intertidal area) were *M. gigas* beds in 2005. Four years later in 2009, wild *M. gigas* accounted for 13.9 kt fresh mass (FM) oyster, whereas cultured *M. gigas* accounted for 6.71 kt FM oyster (Jiang et al., 2019). The specific intertidal *M. gigas* reef, where we collected the specimens for our study, likely originated from a scientific experiment conducted in 2001 (Diederich, 2005).

Here, we aimed (a) to determine the short-term uptake and processing of bacterioplankton by *M. gigas* and its associated fauna in summer and autumn, and (b) to quantify differences in the metabolic activity of intertidal macrozoobenthos in the same seasons.

## Materials and methods

### Experimental design

For this study, *ex-situ* (n = 6) experiments and the corresponding controls were conducted in summer (July) and autumn (September – October) 2020. The controls included the sampling of specimens with natural abundance stable isotopic composition (control 1; n = 6). They were also used to correct for the exchange of noncovalently bound H (control 2; n = 3), and to correct for bacterial respiration and nutrient release and/or uptake (control 3; n = 6).

### Main treatment

Immediately prior to the incubation experiments, *M. gigas* with its epibiont H. *panicea* (hereinafter referred to as ‘*H. panicea*/*M. gigas* combination’) were collected by hand at an *M. gigas* reef in the eastern part of the Eastern Scheldt (51.489°N, 4.058°E; Fig. 1) during low tide and transported to the mesocosm facilities of NIOZ-Yerseke.

**Figure 1.**
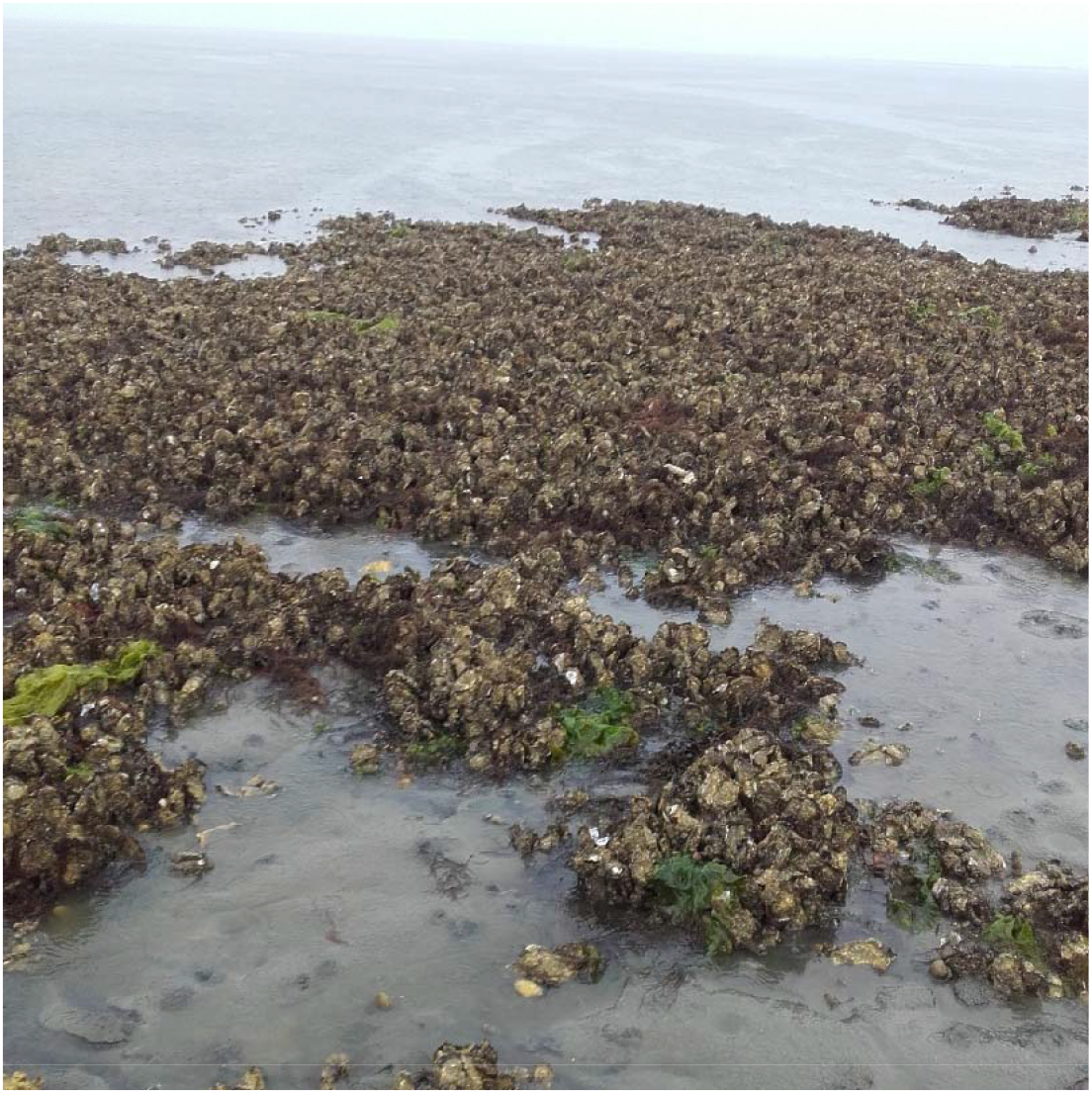
Photo from the intertidal Pacific oyster reef in the Eastern Scheldt where *Magallana gigas* specimens including their associated fauna were collected for the *ex-situ* experiments.

In a temperature controlled-room (temperature *T*_summer_: 18.7°C, *T*_autumn_: 12.6°C) of the mesocosm facility, *H. panicea*-*M. gigas* combinations were placed inside large incubation chambers that were fitted with magnetic stirrers, lids, and sampling ports (chamber volume: 8.5 L; (Stratmann et al., 2025b)). Half of each chamber was filled with pre-filtered seawater (2 μm filter pore size; *S*_summer_: 34.6, *S*_autumn_: 23.4) from the Eastern Scheldt, before 85 ml (summer 2020) and 213 ml (autumn 2020), respectively, deuterium oxide (99.9%) were added, leading to 1% (v/v) and 2.5% (v/v) ^2^H-enrichment. The difference in ^2^H-enrichments was necessary, because nanoscale secondary ion mass spectrometry (NanoSIMS) measurements of *H. panicea* specimens from the autumn 2020 experiments were planned which required a higher ^2^H incorporation. These enrichment differences did affect the results and the data were standardized such that the seasonal data are directly comparable.

After placing the chambers in a water bath, the chambers were filled completely with pre-filtered seawater and closed airtight with lids that had optical oxygen sensors (FireStingO_2_, PyroScience GmbH, Germany) inserted. Duplicate water samples for (^13^C-)DIC and inorganic nutrients (nitrite, ammonia, nitrate, dissolved silicate DSi) were taken through a sampling port (T_-1_), and 3.50 ml freshly prepared ^13^C and ^15^N-enriched concentrated labelled substrate bacteria (summer 2020: no data; autumn 2020: 28.5 at% ^13^C/48.7 at% ^15^N; 10.5 mg ^13^C, 3.41 mg ^15^N) was added. After taking another set of duplicate water samples (T_0_), the sampled water from the incubation chambers was replaced by pre-filtered seawater, and the sampling ports were closed airtight. In the dark, O_2_ consumption inside the chambers was recorded continuously every 10 sec as air saturation (%) of seawater for 4 h, while the water was kept homogenously mixed by stirring with the magnetic stirrers. After terminating the O_2_ measurements, another set of duplicate water samples was taken (T_1_), and the water inside the chambers was aerated via aquarium air pumps until the end of the ‘incubation period’. Further duplicate water samples were taken 6 (T_2_), 8 (T_3_), 10 (T_4_), and 12 h (T_5_) after the begin of the experiments.

Water samples were retrieved for (^13^C-)DIC measurements at T_-1_, T_0_, and T_1_. Sample water was stored in 10 ml headspace vials at 4°C and fixed with 10 μl saturated HgCl_2_. Inorganic nutrient samples (T_-1_, T_0_, T_1_, T_2_, T_3_, T_4_, T_5_) were filtered through 0.45 µm filters into 6 ml vials and stored frozen (-20ºC; nitrite, ammonia, nitrate) or at 4ºC (DSi).

After T_5_, the experiment was terminated by removing the ‘*H. panicea*/*M. gigas* combination’ from the incubation chambers. The specimens were washed in pre-filtered seawater and *H. panicea* specimens were scrapped off the *M. gigas* shells. All epifauna of *H. panicea* and fauna living inside its tissue were handpicked and frozen (-20°C), wet mass (WM) of the *H. panicea* specimens was determined, and the specimens were frozen (-20°C). All epifauna growing on *M. gigas* shells were removed and frozen (-20°C), before the shells were opened with an oyster knife. The flesh was removed, its WM was measured, and it was frozen (-20°C), while the shells were discarded.

### Control 1 (Natural abundance stable isotopic composition)

Six *H. panicea* and six *M. gigas* specimens were collected near the intertidal *M. gigas* reef per season. In the lab, *H. panicea* and *M. gigas* specimens were sampled and preserved as described in “Main treatment”.

### Control 2 (Correction for exchange of noncovalently bound H)

To correct for the exchange of noncovalently bound H with other noncovalently bound H (Schimmelmann et al., 2001; Sessions et al., 2004; Lis et al., 2006), the incorporation of ^2^H in dead *H. panicea* and dead *M. gigas* was investigated as suggested in (Stratmann et al., 2025b). For this purpose, three *H. panicea* and three *M. gigas* specimens per season were collected at the intertidal *M. gigas* reef, and transported to the lab. In the lab, each *H. panicea* specimen was weighed to determine WM; the *M. gigas* specimens were opened with an oyster knife, the flesh removed, and weighed. Each specimen was transferred to a separate sampling container which was filled with 4% borax-buffered formaldehyde until each specimen was fully submerged. After incubating in formaldehyde for 9 h, the formaldehyde was discarded and the *H. panicea* and *M. gigas* specimens were transferred to small incubation chambers equipped with magnetic stirrers and lids (chamber volume: 1.4 l; (Stratmann et al., 2024, 2025b, 2025a)). These chambers were filled with pre-filtered (2 μm pore size) seawater (salinity *S*_summer_: 35.1, *S*_autumn_: 22.7) from the Eastern Scheldt and 14 ml deuterated water (99.9%; equivalent to 1% (v/v) ^2^H-enrichment). The incubation chambers were closed, the magnetic stirrers inside the chambers started stirring, and the dead specimens incubated in a water bath, in the dark for 12 h. Afterwards, the specimens were frozen (-20°C).

### Control 3 (Correction for bacteria contribution to fluxes)

To correct for the contribution of bacteria to fluxes and O_2_ consumption, freshly prepared ^13^C and ^15^N-enriched concentrated labelled substrate bacteria were incubated in control 3. For these experiments, small incubation chambers were filled with pre-filtered seawater (2 μm filter pore size; *S*_summer_: 35.1, *S*_autumn_: 23.2) from the Eastern Scheldt, and placed in a water bath. Duplicate water samples were taken for inorganic nutrients (nitrite, ammonia, nitrate, DSi) and (^13^C-)DIC measurements at T_-1_, before 0.59 ml concentrated labelled substrate bacteria (summer 2020: no data; autumn 2020: 27.4 at% ^13^C/46.2 at% ^15^N; 0.71 µg ^13^C, 0.26 µg ^15^N) were added. Another single water sample was taken for inorganic nutrients and (^13^C-)DIC measurements (T_0_) and the missing water was replaced with pre-filtered seawater before the chambers were closed airtight with lids in which optical O_2_ sensors were inserted. The air saturation (%) of seawater was recorded continuously in the dark for 4 h in the dark, while the water was stirred with the magnetic stirrers. At the end of the O_2_ measurements, a single water sample was taken for inorganic nutrients and (^13^C-)DIC at T_1_. Further duplicate water samples for inorganic nutrients were taken 6 h after the begin of the incubations (T_2_), 8 h (T_3_), 10 h (T_4_), and 12 h (T_5_) later. Water samples for inorganic nutrients and (^13^C-)DIC analyses were processed and preserved as described in “Main treatment”.

### Sample processing

#### Fauna samples

Epifauna found associated with the ‘*H. panicea*/*M. gigas* combination’ was counted and identified to the finest taxonomic level possible (i.e., mostly species/genus level for adult specimens, higher taxonomic level for juveniles) under a stereomicroscope using (Hayward and Ryland, 1995; Bos et al., 2016) as references. Afterwards, every epifaunal specimen was freeze dried, weighed with (i.e., dry mass DM with shell) and without shell (i.e., DM), and ground to powder using mortar and pestle. *H. panicea* and *M. gigas* specimens were freeze dried, weighed to determine DM, and ground to powder. Org. C content/δ^13^C, TN content/δ^15^N, and H content/δ^2^H of every specimen were measured with an elemental analyzer coupled to an isotope ratio mass spectrometer (EA-IRMS) as described in detail in (Stratmann et al., 2025b). C, N, and H stable isotope values were reported in δ notation relative to Vienna Pee Dee Belemnite (VPDB) for δ^13^C, relative to air for δ^15^N (Fry, 2006), and relative to Standard Mean Ocean Water normalized to Standard Light Antarctic Precipitation (VSMOW-SLAP) for δ^2^H (Coplen, 1995).

#### Inorganic nutrient and dissolved inorganic carbon analyses

Inorganic nutrients (ammonia, nitrite, nitrate, DSi) concentrations (µmol l^-1^) were quantified with a SEAL QuAAtro analyzer (Bran+Luebbe, Germany), after warming the samples to room temperature for 24 h. DIC concentrations (µmol l^-1^) were measured with an Apollo SciTech DIC analyzer (AS-C3; Apollo SciTech, USA) as described in (Moodley et al., 2000). The isotopic composition of DIC was measured by converting DIC into CO_2_ as described in (Stratmann et al., 2025b) and by measuring its δ^13^Cvalue with Thermo Delta V continuous flow IRMS.

### Data analysis

#### Inorganic nutrient and dissolved inorganic carbon flux calculations

Biomass-corrected fluxes *Flux*_*(MT, parameter)*_ (nitrate, nitrite, ammonia, DSi, ^13^C-DIC, DIC; μmol mmol C^-1^ d^-1^) of the main treatment were calculated as follows:

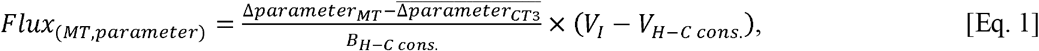

where Δ*parameter* is the change of the concentration (μmol l^-1^) of a specific parameter over time (T_0_ to T_5_; d) determined via robust regression analyses (α = 0.05).*B*_*H-Ccons*_. is the biomass (mmol C) of the ‘*H. panicea*/*M. gigas* combination’ and any epifauna present. *V*_*I*_ is the volume of the incubation chamber (= 8,500 ml) and *V*_*H-C cons*._ is the volume of the *H. panicea*/*M. gigas* combination (ml) which was calculated as:

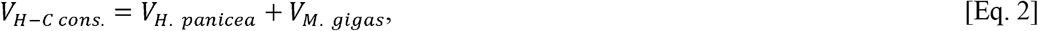

where

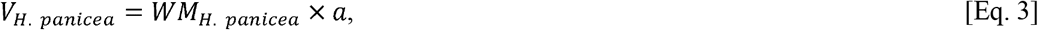

*V*_*H*. *panicea*_ is the volume (ml) of the individual *H. panicea, WM*_*H*. *panicea*_ is the wet mass (g) of *H. panicea*, and *a* (= 0.73 ml g^-1^) is the WM:V conversion factor for *H. panicea* from the Eastern Scheldt (Stratmann et al., 2025b).

*V*_*H-C cons*._ as volume (ml) of *M. gigas* is calculated following (Ren, 2001) as:

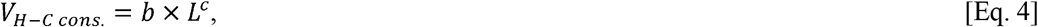

with *L* being the length (cm) of the shell of *M. gigas. b* is 0.38 ml cm^-1^ and c is 2.15.

### Oxygen consumption calculations

Oxygen consumption 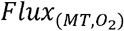 (μmol O_2_ mmol C d^-1^) in *ex-situ* experiments was calculated as described in (Stratmann et al., 2024, 2025b). Air saturation (%) of seawater was converted to absolute O_2_ concentrations (mg O_2_ l^-1^) following (Weiss, 1970) using the *gas_O2sat* function in the ‘marelac’ package (vs. 2.1.11) (Soetaert and Petzoldt 2008) in *R* (R-Core Team, 2025). Afterwards, the decline of O_2_ concentrations over time (ΔO_2_, μmol O_2_ l^-1^ d^-1^) was determined by linear regression (α = 0.05) and blank-corrected and biomass-corrected 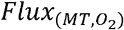 was calculated as:

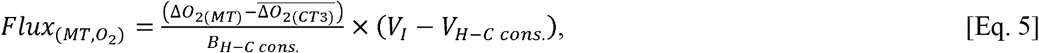

where 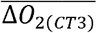 is the average decline of O_2_ concentrations over time in the control 3 incubations of a specific season (summer 2020, autumn 2020).

### Calculations of bulk ^13^C, ^15^N, and ^2^H uptake

The bulk uptake of ^13^C, ^15^N, and ^2^H by fauna was calculated as described in (Stratmann et al., 2025b):

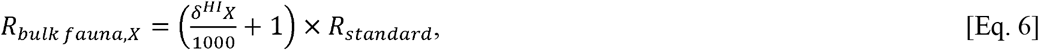

where *X* is C, N, or H, ^HI^X is the heavy stable isotope of C, N, or H (i.e., ^13^C, ^15^N, ^2^H), and *R*_*standard*_ is 0.01118 (for C), 0.0036765 (for N), and 0.00015576 (for H) (Fry, 2006). The fraction of the specific heavy stable isotope *F*_*bulk fauna,X*_ is:

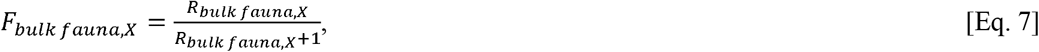

and the bulk uptake of ^13^C, ^15^N, and ^2^H by the individual specimens *I*_*bulk fauna,X*_ (mg ^13^C ind.^-1^, mg ^15^N ind.^-1^, mg ^2^H ind.^-1^) is:

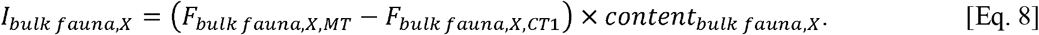

Here, *content*_*bulk fauna,X*_ is the concentration (mg ind.^-1^) of C, N, and H of the specific specimen.

Biomass-corrected ^13^C, ^15^N, and ^2^H uptake of fauna *I*_*bulk fauna,X,bio-cor*_ (mg mg C^-1^) is:

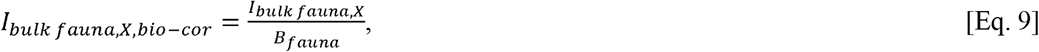

with *B*_*fauna*_ being the biomass of the individual specimen (mg C ind.^-1^).

We corrected for the exchange of noncovalently bound H in bulk *H. panicea* and *M. gigas* tissue (Δ*I*_*bulk HP&MG,H*_) as follows:

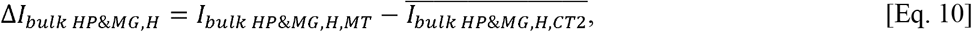

Whereupon 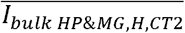 data are presented in Table 1.

**Table 1.** Bulk uptake of ^2^H (×10^-5^ mg ^2^H mg C^-1^) by *Magallana gigas* and *Halichondria panicea* in control 2 during summer and autumn 2020. Data are presented as mean ± SD (median; 1^st^ – 3^rd^ quartile).

| Species | Summer 2020 | Autumn 2020 |
| --- | --- | --- |
| *Halichondria panicea* | 7.498 ± 10.37<br>(2.690, 0.643 – 8.518) | 1.206 ± 1.564<br>(0.643, 0.322 – 1.808) |
| *Magallana gigas* | 3.333 ± 1.460<br>(3.701, 2.712 – 4.137) | 2.510 ± 2.181<br>(3.590, 1.795 – 3.766) |

No direct measurements of abiotic uptake of ^2^H by epifauna species was investigated in this study. Therefore, we corrected for noncovalently bound H exchange in epifauna species by subtracting 10% of the bulk ^2^H uptake, because (Stratmann et al., 2025b) found that living shallow-water sponges took up ten times more ^2^H than dead sponges from control 2:

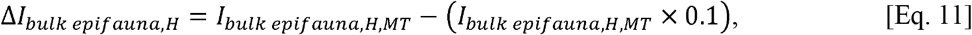

Biomass-corrected tracer H uptake of fauna *I*_*bulk fauna,tracer H,bio-cor*_ (mg tracer H mg C^-1^) was calculated as:

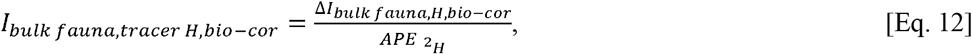

where 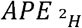 is the atom% excess of ^2^H-enriched seawater (i.e., 1 atom% for experiments in summer 2020; 2.5 atom% for experiments in summer 2020).

The ratios of ^13^C uptake/^15^N uptake 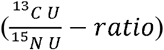, ^13^C uptake/tracer H uptake 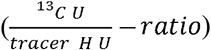,and ^15^N uptake/tracer H uptake 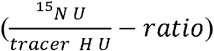 were calculated as:

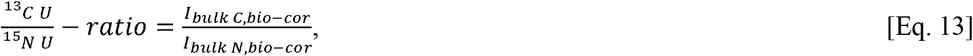

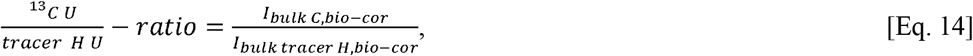

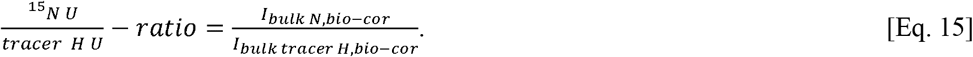

The latter two ratios provide information about how dependent the fauna is on bacterioplankton C 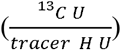 and 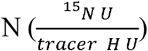.

### Statistical data analysis

It was tested whether *Flux*_(*MT*_,_*parameter*)_ differed significantly from 0 μmol mmol C^-1^ d^-1^ by performing two-tailed 1-sided Student’s t-tests (*t*; α = 0.05) in *R*, after normality was checked by Shapiro-Wilk normality tests. Non-normally distributed flux data were assessed by 1-sample Wilcoxon tests (*W*; α = 0.05) in *R*. Subsequently, a seasonal difference (summer vs. autumn) in *Flux*_(*MT*_,_*parameter*)_was tested with an unpaired two-samples t-test (α = 0.05) and a Wilcoxon Rank-Sum Test (α = 0.05) in *R*.

Differences in bulk ^13^C, ^15^N, and ^2^H uptake (*I*_*bulk fauna,X,bio-cor*_) among sampling seasons (summer 2020, autumn 2020) were tested with Welch’s t-tests (*t*; α = 0.05) in case of normally distributed data and with Mann-Whitney-Wilcoxon test (*V*, α = 0.05) for non-normally distributed data, after normality was assessed by the Shapiro-Wilk normality test.

Data are presented as mean ± SD (median).

## Results

### Epifauna composition

The *H. panicea* and *M. gigas* specimens selected for the *ex-situ* experiments hosted in total 24 different epifauna taxa (Table 2). In summer 2020, these taxa belonged to the phyla Arthropoda, Echinodermata, Mollusca, and Porifera, whereas in autumn 2020, the phyla Annelida, Arthropoda, Echinodermata, Mollusca, Nemertea, Platyhelminthes, and Porifera were present.

**Table 2.** Macrozoobenthos epi- and endofauna of *Magallana gigas* and *Halichondria (Halichondria) panicea* present (+) or absent (-) in summer and autumn 2020.

| Taxon | Summer 2020 | Autumn 2020 |
| --- | --- | --- |
| *Amphiura* sp. | + | - |
| *Carcinus maenas* | + | + |
| Caridea | - | + |
| *Crepidula fornicata* | - | + |
| *Eriocheir sinensis* | + | + |
| *Haliclona* (*Haliclona*) *oculata* | + | - |
| *Hymeniacidon perlevis* | + | + |
| *Littorina littorea* | + | + |
| Monoplacophora | - | + |
| *Mytilus edulis* | + | - |
| Nemertea | - | + |
| Nereididae | - | + |
| *Ophiothrix fragilis* | + | - |
| Ophiuroidea | + | + |
| Patellidae | + | - |
| *Pisidia longicornis* | - | + |
| *Polybius depurator* | + | + |
| Polychaeta (undefined) | + | + |
| Pycnogonida | + | + |
| *Rhithropanopeus harrisii* | - | + |
| Turbellaria | - | + |
| *Urosalpinx cinerea* | + | + |

### Fluxes of inorganic nutrients, dissolved inorganic carbon, and oxygen

*Ex-situ* fluxes of inorganic nutrients varied substantially between sampling seasons (Fig. 2, Table 3): Nitrate was consumed in summer 2020, but released in autumn 2020, whereas nitrite and ammonia were released during both sampling seasons. Dissolved silicate was consumed and total DIC was produced, whereupon ^13^C-DIC was consumed in summer 2020, but released in autumn 2020. Oxygen was always consumed at significant rates (*t*(5)_summer_ = -7.31, *p* = 0.001; *t*(5)_autumn_ = -6.06, *p* = 0.002), but 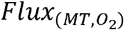 did not differ between seasons (*t*(8.99) = 0.75, *p* = 0.48).

**Table 3.** Results of Student’s t-test (α = 0.05) and Wilcoxon test (α = 0.05) to assess whether the significant fluxes (μmol mmol C^-1^ d^-1^; Supplementary Table 6) of inorganic nutrients, DIC, and ^13^C-DIC differed significantly between seasons in 2020. Symbols: \**p*-value≤0.05, □*p*-value≤0.01, □*p*-value≤0.001.

| Parameter | t-value, W-value | Df | p-value |
| --- | --- | --- | --- |
| $\text{NO}_2^-$ | -2.823 | 5.87 | 0.031* |
| $\text{NH}_4^+$ | 31 | | 0.041* |
| $^{13}\text{C}$-DIC | -9.465 | 5.25 | 0.0002□ |

**Figure 2.**
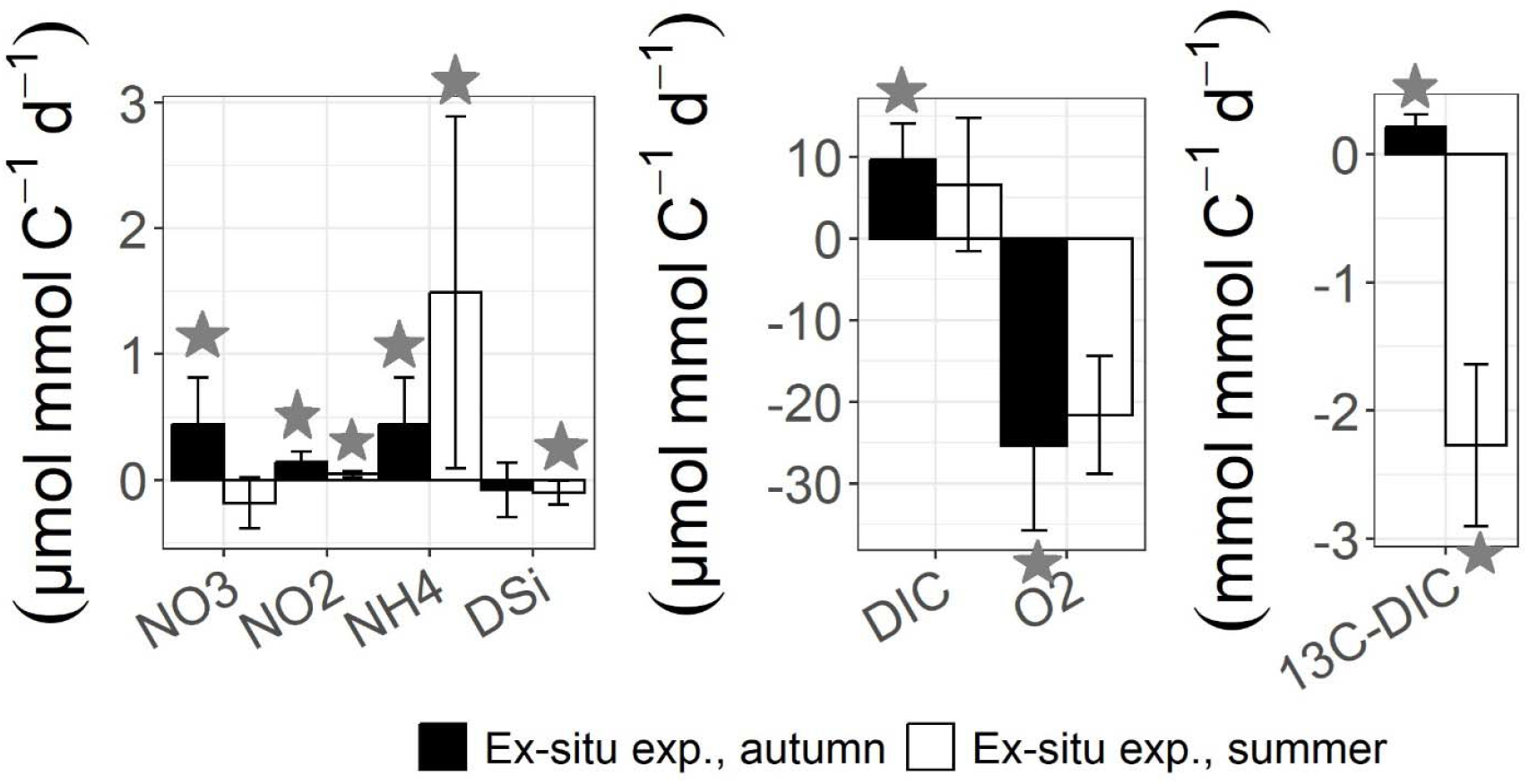
Fluxes (*Flux*, μmol mmol C^-1^ d^-1^, mmol mmol C^-1^ d^-1^) of nitrate (NO_3_ ^-^), nitrite (NO_2_ ^-^), ammonium (NH_4_ ^+^), dissolved silicate (DSi), DIC, ^13^C-DIC, and oxygen (O_2_) of the ‘*Halichondria panicea*/*Magallana gigas* combination’ measured in summer and autumn 2020. Negative fluxes signify that a specific compound was taken up by the ‘*H. panicea*/*M. gigas* combination’, whereas a positive flux corresponds to the release of the compound. A star indicates that this average flux was significant (see also Supplementary Table 6). Error bars represent 1 SD.

### Biomass-specific uptake of C, N, and H

During the experiments, the macrozoobenthos (i.e., epifauna, *H. panicea*, and *M. gigas*) consumed 3.00 ± 1.72 (2.88) µg ^13^C mg C^-1^, 1.43 ± 0.71 (1.39) µg ^15^N mg C^-1^, and 1.05 ± 1.41 (0.53) µg tracer H mg C^-1^ in summer 2020, and 8.73 ± 1.64 (8.66) µg ^13^C mg C^-1^, 3.75 ± 0.68 (3.65) µg ^15^N mg C^-1^, and 1.39 ± 1.19 (1.09) µg tracer H mg C^-1^ in autumn of the same year (Fig. 3). This difference in uptake of ^13^C, ^15^N, and tracer H between the two seasons was significant for ^13^C (*t*(9.97) = -5.91, *p*-value = 0.0002) and ^15^N (*t*(9.98) = -5.84, *p*-value = 0.0002) uptake, but not for tracer H uptake (*W* = 13, *p*-value = 0.48). Even when all species were excluded that were only present in one of the two seasons (i.e., Nereididae, Ophiuroidea, *O. fragilis, Amphiura* sp., *Haliclona (Haliclona) oculata*, Caridea, *Crepidula fornicata, Rhithropanopeus harrisii*, Monoplacophora, *Pisidia longicornis*, Platyhelminthes), the difference in uptake of ^13^C, ^15^N, and tracer H between the two seasons was significant for ^13^C (*t*(9.04) = - 3.54, *p*-value = 0.006) and ^15^N (*t*(9.30) = -3.48, *p*-value = 0.007) uptake, but not for ^2^H uptake (*W* = 14, *p*-value = 0.59).

**Figure 3.**
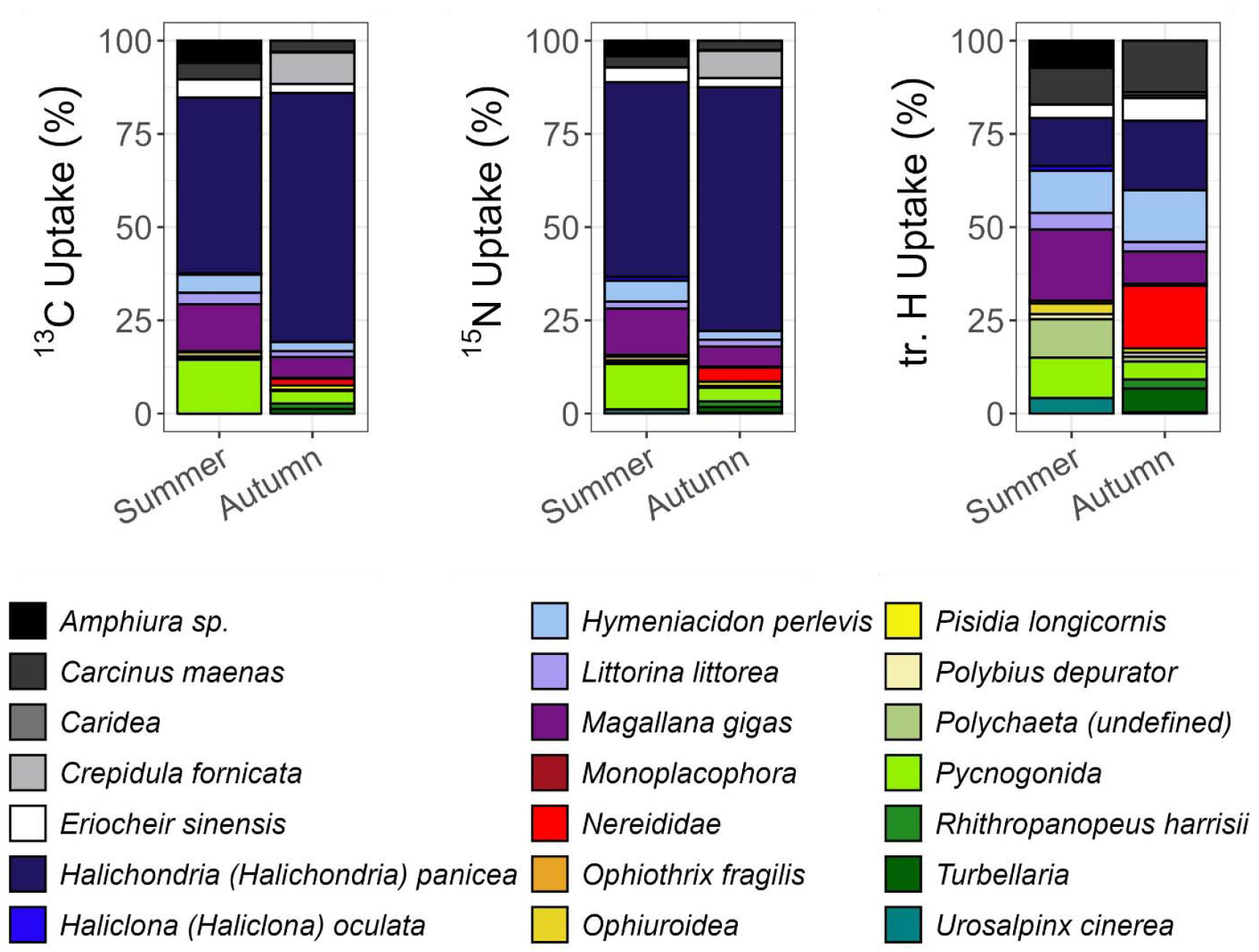
Relative biomass-specific uptake of ^13^C, ^15^N, and tracer H (% total biomass-specific uptake) by fauna in summer and autumn 2020.

Most total biomass-specific ^13^C was taken up by *H. panicea, Eriocheir sinensis*, and *Carcinus maenas* in summer, and by *H. panicea, C. fornicata*, and *R. harrisii* in autumn 2020, whereas most total biomass-specific ^15^N was incorporated into *H. panicea, E. sinensis*, and *Hymeniacidon perlevis* in summer and into *H. panicea, C. fornicata*, and *R. harrisii* in autumn 2020. In comparison, most relative biomass-specific ^13^C and ^15^N were taken up by *H. panicea*, Pycnogonida, and *Magallana gigas* in summer, and by *H. panicea, C. fornicata*, and *M. gigas* in autumn 2020 (Fig. 3).

Total biomass-specific tracer H uptake was dominated by *C. maenas, E. sinensis*, and *H. perlevis* in summer and by *H. perlevis, R. harrisii*, and *Littorina littorea* in autumn 2020. In comparison, most relative biomass-specific tracer H was incorporated into *M. gigas, H. panicea*, and *H. perlevis* in summer, and in *H. panicea*, Nereididae, and *H. perlevis* in autumn 2020 (Fig. 3).

The — -ratios of macrozoobenthos ranged from 0.90 (*H. oculata*) to 3.80 ± 0.11 (3.82) (*L. littorea*) in summer 2020 (Fig. 4, upper panel) and from 0.90 ± 0.44 (0.74) (Nereididae) to 2.97 ± 0.26 (2.94) (*C. maenas*) in autumn 2020 (Fig. 4, lower panel).

**Figure 4.**
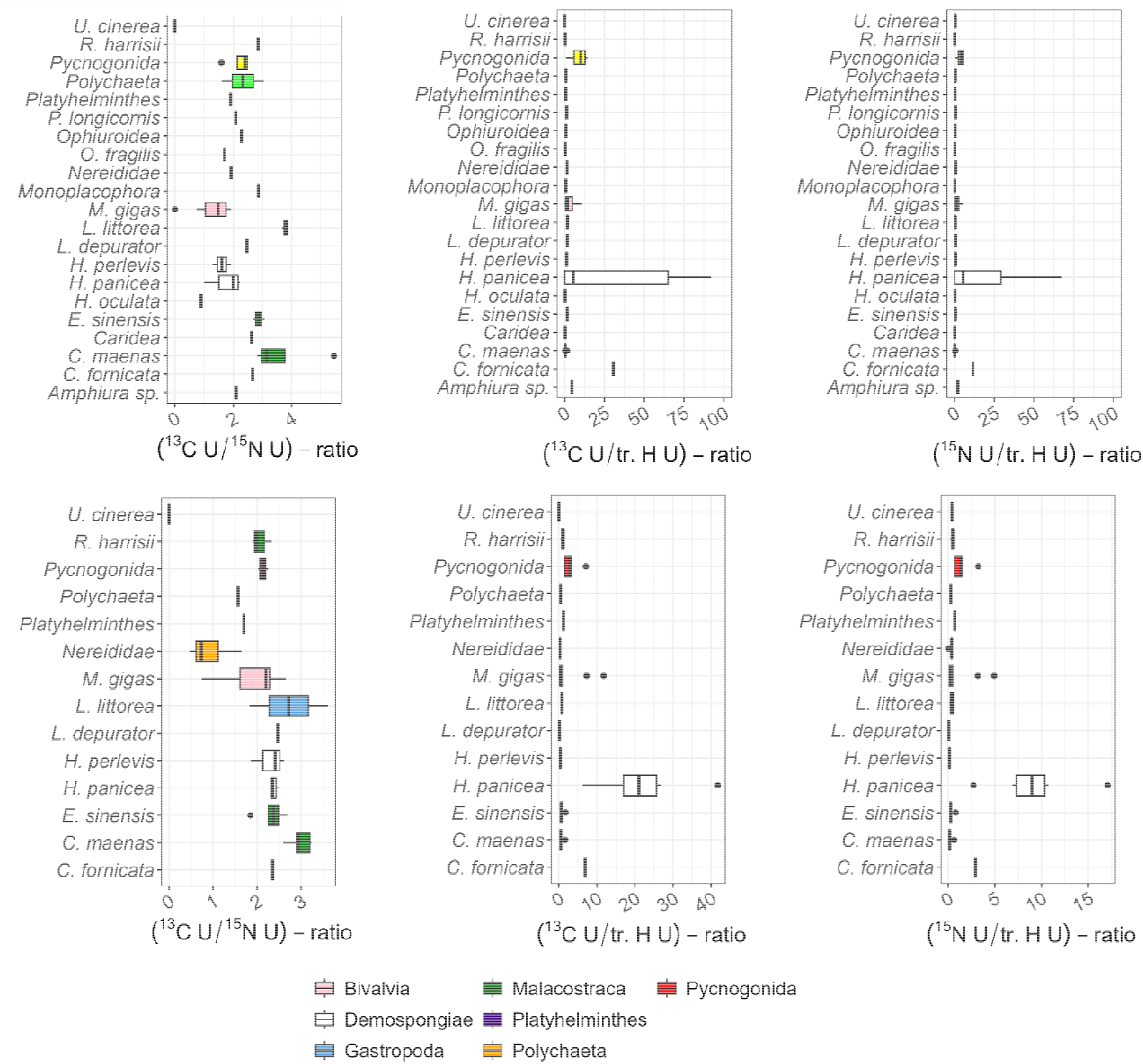
Ratios of —, —, and of — macrozoobenthos species present in (upper panel) summer and in (lower panel) autumn 2020. For presentation reasons, the x-axis of the panels — and — of summer 2020 were limited to 100 excluding one *H. panicea* outlier.

In summer 2020, *H. panicea* had the highest 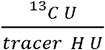 and 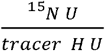 ratios of all studied macrozoobenthos 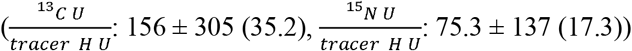 and the lowest 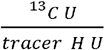and 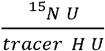-ratios were found in *H. oculata* 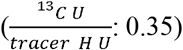 and *R. harrisii* 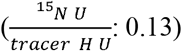, respectively (Fig. 4, upper panel, Fig. 5).

**Figure 5.**
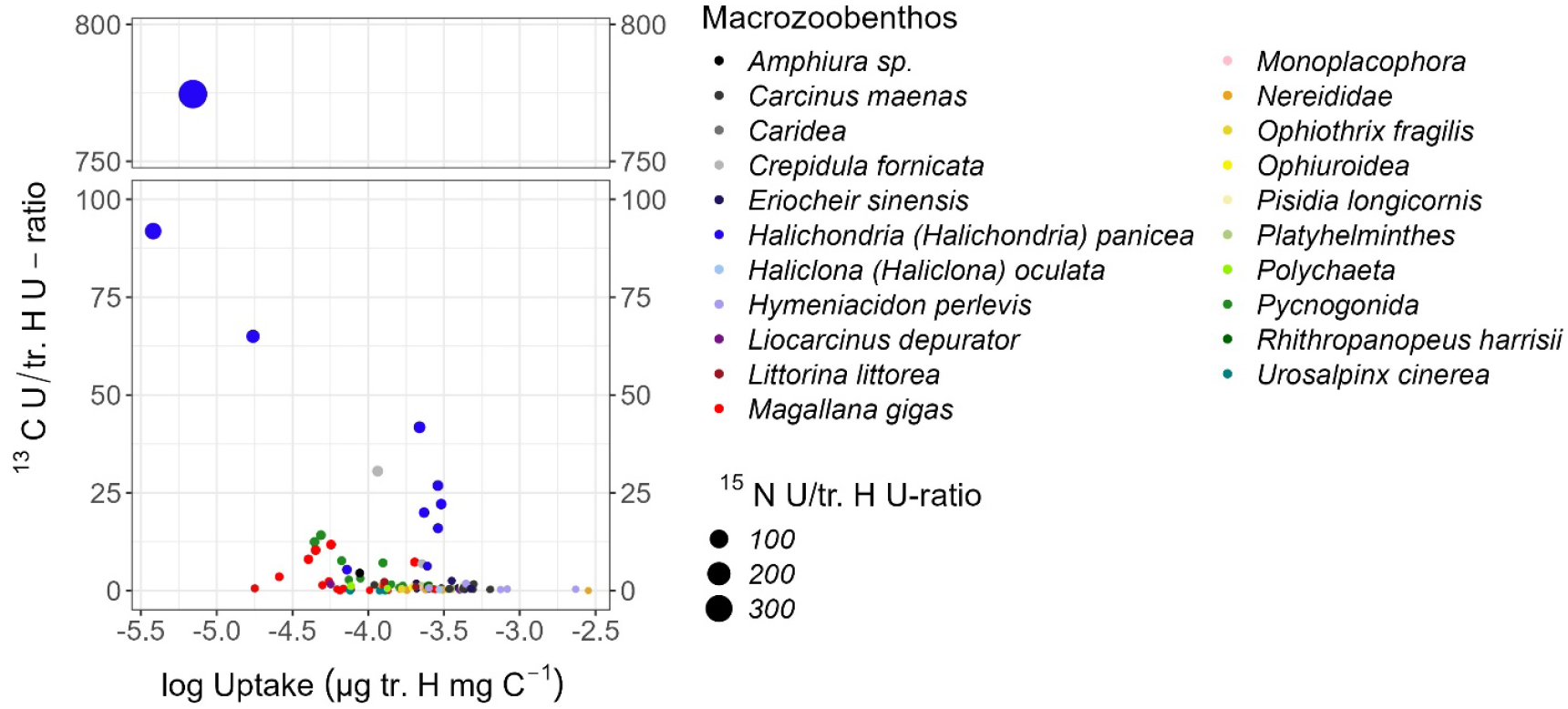
Biomass-specific uptake of tracer H (µg tracer H mg C^-1^) versus 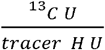 – and 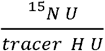 -ratios for individual macrozoobenthos specimens. Note that the biomass-specific uptake rate of tracer H is presented as decadal logarithm of respective uptake rate.

In comparison, in autumn 2020, the 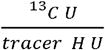 – and 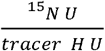 -ratios of macrozoobenthos ranged from *P. depurator* 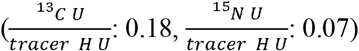 to *H. panicea* 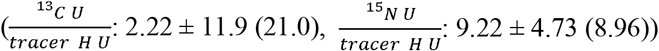 (Fig. 4, lower panel).

Bivalvia was the class with the lowest 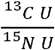-ratio in summer (1.29 ± 0.65 (1.48)) and Polychaeta was the class with the lowest 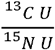 -ratios in autumn 2020 (0.99 ± 0.47 (0.80)), whereas Malacostraca was the class with the highest 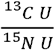-ratios in summer (3.09 ± 0.89 (2.86)) and autumn 2020 (2.51 ± 0.43 (2.47)).

In summer 2020, the class with the highest 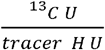 – and 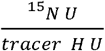 -ratio was Demospongiae 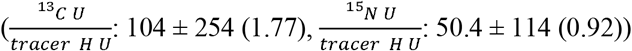,whereas the lowest ratios were detected in Platyhelminthes 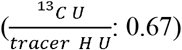 and Monoplacophora 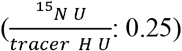. In autumn 2020, classes with the lowest 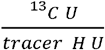 – and 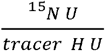 -ratios were Polychaeta (0.34 ± 0.19 (0.35)) and Malacostraca (0.32 ± 0.23 (0.24)), whereas Demospongiae had the highest - — and —-ratios (—: 14.9± 14.4 (16.0), —: 6.20 ± 5.88 (6.89)).

When feeding types were considered, the mean —-ratios increased from omnivores, filter and suspension feeders, carnivores, to deposit feeders and finally herbivores (Fig. 6, left panel). The feeding types with the highest mean —- and —-ratios were filter and suspension feeders, followed by carnivores, deposit feeders, herbivores and omnivores (Fig. 6, middle and bottom panel).

**Figure 6.**
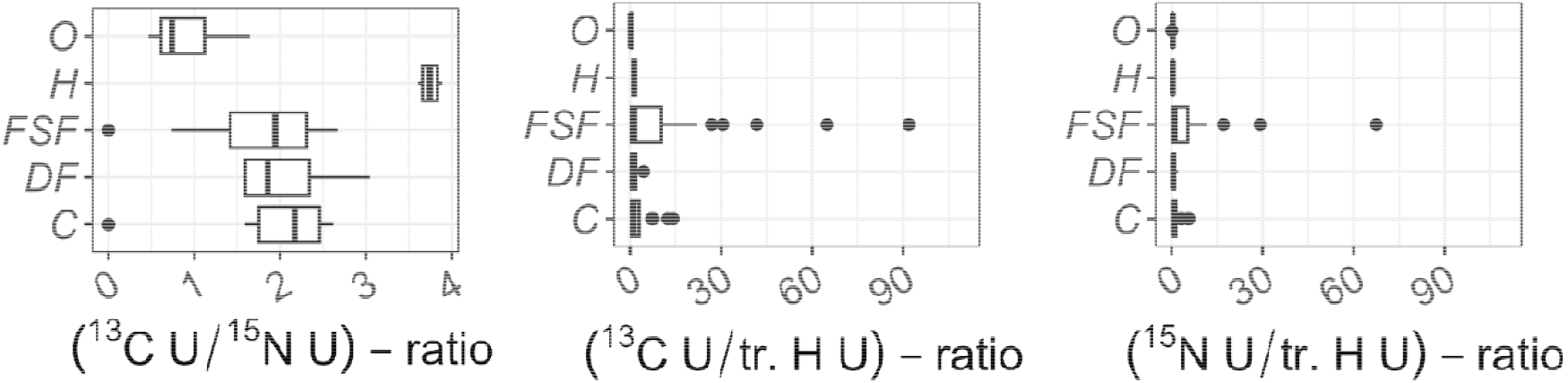
Ratios of —, —, —and of adult macrozoobenthos belonging to different feeding types, excluding very high data points. Species were classified in feeding types based on published literature (Table S9). Abbreviations: C = carnivore, DF = deposit feeder, FSF = filter and suspension feeder, H = herbivore, O = omnivores.

## Discussion

### Short-term benthic community responses to pelagic bacteria

Over a period of 12 h, most ^13^C and ^15^N per biomass was taken up by filter and suspension feeders (i.e., *H. panicea, C. fornicata, H. perlevis*), predators (i.e., *C. maenas*), and omnivores (i.e., *E. sinensis, R. harrisii*) (Fig. 7).

**Figure 7.**
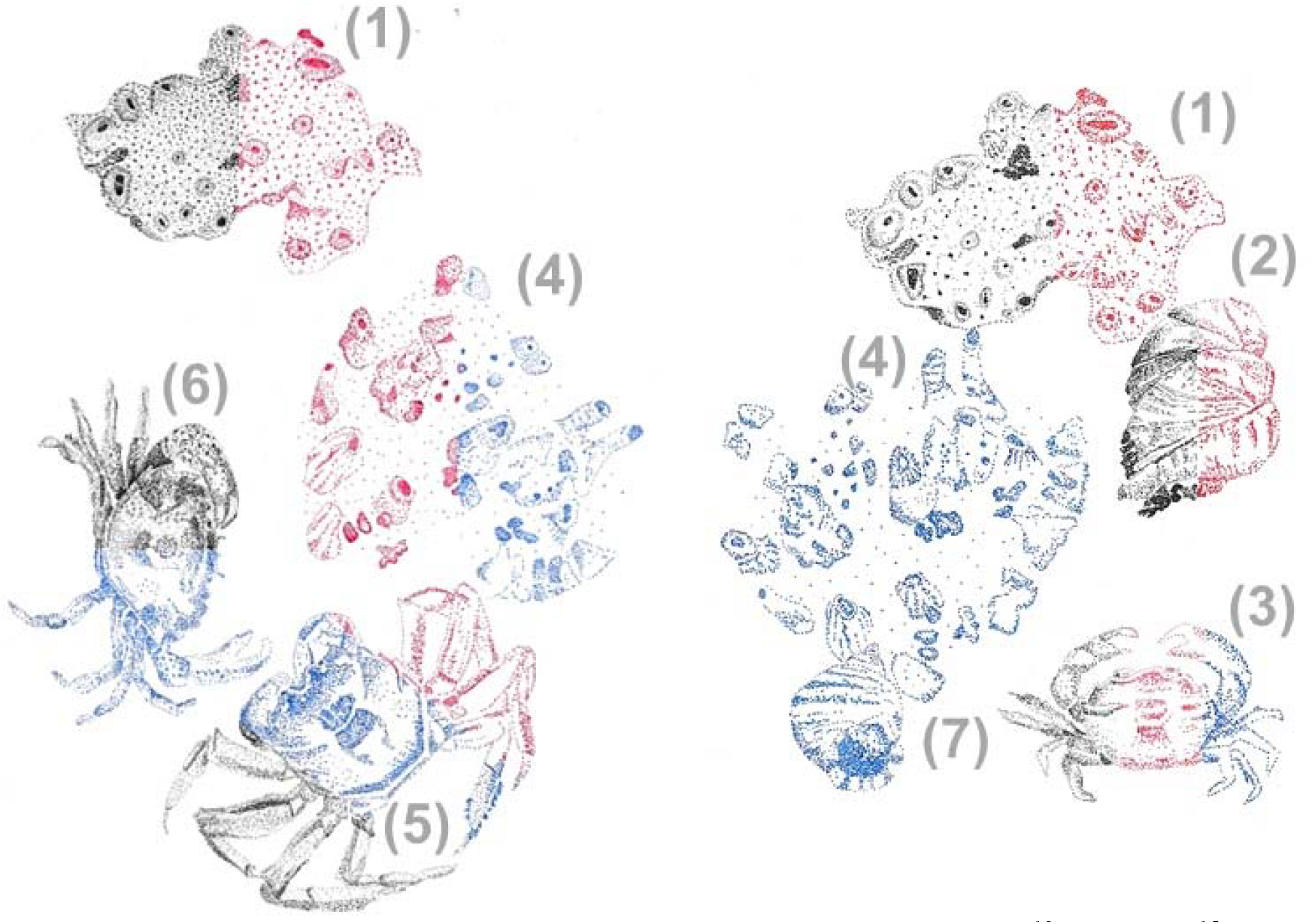
Drawings of the three macrozoobenthos species that took up most ^13^C (black), ^15^N (red) and ^2^H (blue) on a biomass (mg C) level per season (left panel) summer vs. (right panel) autumn 2020. A multicolor drawing of a species indicates that this species belongs the three species with the highest uptake for multiple stable isotopes. The species presented are: (1) *Halichondria* (*Halichondria*) *panicea*, (2) *Crepidula fornicata*, (3) *Rhithropanopeus harrisii*, (4) *Hymeniacidon perlevis*, (5) *Eriocheir sinensis*, (6) *Carcinus maenas*, (7) *Littorina littorea*. Illustrations by Tanja Stratmann.

The filter-feeding ***H. panicea*** is the dominant intertidal sponge in the Eastern Scheldt (de Kluijver and Leewis, 1994), where it grows attached to macroalgae, *M. gigas*, roof tiles, and bricks (Stratmann et al., 2025b). This species filters phytoplankton and free-living bacteria out of the water column (Riisgård et al., 2016) with rates of 27.0 to 28.4 ml water g^-1^ DM sponge min^-1^ (Riisgård et al., 1993; Thomassen and Riisgård, 1995). In this study, *H. panicea* took up approximately three times more bacteria-derived ^13^C in autumn than in summer 2020. Assuming that the ratio of phyto-vs. bacterioplankton biomass in the Eastern Scheldt changes seasonally similar to the ratio in the Kerteminde Fjord (Baltic Sea) (Lüskow et al., 2019), *H. panicea* is more exposed to bacterioplankton in autumn than to phytoplankton and therefore it is potentially adapted to feeding more upon free-living bacteria in autumn and phytoplankton in summer. Another explanation could be that *H. panicea* is more food-limited in autumn than in summer, and therefore retains more bacteriaplankton-derived ^13^C in autumn. (Stratmann et al., 2025b) studied the uptake of ^13^C and ^15^N-labeled substrate bacteria by *H. panicea* from the Eastern Scheldt in summer 2019 and assessed a potential food limitation of this species. However, the results remained inconclusive for July 2019 and the authors did not study specimens in September/October 2019, so at this stage we cannot confirm whether *H. panicea* is indeed more food limited in autumn than in summer.

In comparison, the non-native sponge ***H. perlevis*** which reached the Netherlands via the transport of shellfish in 1951 (Gittenberger et al., 2017) is less abundant in the Eastern Scheldt and was only found at the sampling site in autumn 2020, but not in summer of the same year (Stratmann, own observations). This species is among the taxa with the highest short-term biomass-specific uptake of bacteria-derived ^13^C and ^15^N because it filters bacterioplankton more efficiently than other filter-feeding bivalves. In a laboratory experiment with *H. perlevis* and the mussel *Mytilus galloprovincialis, H. perlevis* took up more bacteria than *M. galloprovincialis* independent of the presence or absence of the mussels (Longo et al., 2016). However, the efficiency with which *H. perlevis* filters the bacterioplankton is also dependent on its community composition. In a different laboratory study *H. perlevis* removed 60% of the initial concentration of the bacterium *Escherichia coli* independent of the initial *E. coli* concentration, whereas the sponge was not able to control or reduce the concentration of the bacterium *Vibrio anguillarum* (Maldonado et al., 2010). Unfortunately, we did not study the community composition of the bacterioplankton cultures we fed to the ‘*H. panicea*/*M. gigas* combination’, so we lack information about which bacteria species were present, but it seems that these were species that *H. perlevis* preferred. The non-native limpet ***C. fornicata***, present in the Netherlands since it was transported to the country with oysters in 1926 (Gittenberger et al., 2017), incorporated 5.09 µg ^13^C mg C^-1^ and 1.98 µg ^15^N mg C^-1^ in its tissue in autumn 2020. This result is intriguing because most studies that compared the diets of *C. fornicata* with the diets of other suspension feeders neglected bacterioplankton as a potential food source. They either determined the filtration, ingestion, and/or retention rates of various phytoplankton and microphytobenthos species (Chaparro et al., 2004; Barillé et al., 2006; Beninger et al., 2007; Harke et al., 2011), or they missed bacterioplankton as input for their stable isotope mixing models (Riera et al., 2002; Decottignies et al., 2007b; Riera, 2007). In a study of *C. fornicata* from the Eastern Scheldt where the limpets use mostly *M. gigas* as substrate (Riera et al., 2002), Riera et al. detected differences in the δ^13^C and δ^15^N signatures of *C. fornicata* and *M. gigas* (Riera et al., 2002). Based on their results, the authors suggested that *C. fornicata* and *M. gigas* ingested and/or assimilated microphytobenthos and suspended particulate organic matter (POM) in different portions. However, they neglected the potential contribution of bacterioplankton to the diet of *C. fornicata* and *M. gigas*. With our study, in which we fed *C. fornicata* and *M. gigas* from the Eastern Scheldt with ^13^C- and ^15^N-enriched bacterioplankton and detected a one order lower ^13^C incorporation into *M. gigas* specimens compared to *C. fornicata* specimens, we can confirm Riera et al.’s hypothesis that competition for food between *M. gigas* and *C. fornicata* is unlikely, because of differences in their diet (Riera et al., 2002). These differences could also be related to *C. fornicata*’s capacity to retain smaller particles better than *M. gigas*: In a laboratory experiment, the retention efficiency of *C. fornicata* increased with the size of the particles from 2 μm to 4 μm equivalent spherical diameter (ESD); particles with a size of >4 μm ESD were retained with 100% efficiency (Barillé et al., 2006). The retention efficiency of *M. gigas* for particles of 3 to 4 μm EDS rose similarly with size, but *M. gigas* only fully retained (i.e., 100%) particles with sizes of >10.4 and 15.7 μm EDS, depending on their chemical composition (Barille et al., 1992).

It might be surprising that crabs were among the taxa that incorporated most bacterioplankton derived ^13^C and ^15^N, but *C. maenas, E. sinensis*, and *R. harrisii* change their diets and feeding types/preferences with developmental stage and/or size: Zoea larvae and post larvae (i.e., megalopa) of ***C. maenas*** are suspension feeders and capture phytoplankton (Harms and Seeger, 1989; Factor and Dexter, 1993; Harms et al., 1994) and small zooplankton (Harms and Seeger, 1989; Harms et al., 1994), whereas juvenile *C. maenas* are predators and predate mostly upon barnacles and to a lesser degree upon gastropods (Rangeley and Thomas, 1987). Adult *C. maenas*, in comparison, prey upon gastropods, barnacles, bivalves, shrimps, polychaetes (Ameyaw-Akumfi and Hughes, 1987; Rangeley and Thomas, 1987; Raffaeffli et al., 1989; Baeta et al., 2006). In this study, all *C. maenas* specimens were juveniles. It is therefore unlikely, that they took up the bacterioplankton directly, but rather that they predated upon sponges present in the incubations, such as *H. panicea, H. perlevis*, or *H. oculata*, and took up the bacterioplankton-derived ^13^C and ^15^N indirectly. It is known that juveniles of *C. maenas* use *M. edulis* beds and lumps as refuge to hide from their epibenthic predators (Thiel and Dernedde, 1994). At intertidal sites in the Eastern Scheldt, they seem to behave similarly and seek refuge in lumps of *M. gigas*. It is also possible that they live associated with *H. panicea* like *C. maneas* at Menai Strait (Irish Sea) (Peattie and Hoare, 1981) and benefit from the protection of the sponge against strong currents, while they predate upon *H. panicea* at the same time.

In their native environment in Asia, the megalopas of ***E. sinensis*** prey upon members of the pelagic food web, such as zooplankton and bacterioplankton, whereas juveniles of *E. sinensis* feed on macrophytes and macrozoobenthos (Jin et al., 2001; Cui et al., 2021). Adult *E. sinensis*, in comparison, consume macrophytes, macrozoobenthos, and forage on fish (Mao et al., 2016). In Europe, to where this species was introduced with ballast water (Gittenberger et al., 2017), adult *E. sinensis* predate upon bivalves, amphipods, gastropods, polychaetes, but they also consume macrophytes and detritus (Wójcik-Fudalewska et al., 2019). Based on our results, we propose that juveniles of *E. sinensis* also predate upon sponges or take up sponge-derived detritus and in this way they are secondary consumers of bacterioplankton-derived ^13^C and ^15^N.

The non-native ***R. harrisii***, which was introduced to the Netherlands before 1874 as biofouling on ship’s hulls (Gittenberger et al., 2017), ingests different items depending on the size of *R. harrisii*. Small *R. harrisii* specimens (<12 mm carapace width) feed detritus, algae, and macrophytes, whereas large *R. harrisii* specimens (>12 mm carapace width) predate upon macrozoobenthos, such as polychaetes and amphipods (Aarnio et al., 2015). Unfortunately, the authors of the study did not investigate the δ^13^C and δ^15^N signature of bacterioplankton as potential food source for small or large *R. harrisii* specimens, so we cannot completely exclude that *R. harrisii* also feeds on bacterioplankton. However, the authors found that small specimens are primary consumers, whereas large specimens are secondary consumers (Aarnio et al., 2015). Hence, we propose that the small *R. harrisii* specimens we detected in the ‘*H. panicea*/*M. gigas* combination’ did indeed filter suspended bacterioplankton out of the water column and incorporated its ^13^C and ^15^N.

### Seasonal differences in C and N cycling

Combining information about ^13^C, ^15^N, and ^2^H uptake with inorganic nutrient and DIC flux rates provides insights into the short-term benthic community response to pelagic bacteria. In **summer 2020**, bacterioplankton was mostly taken up by sponges (i.e., *H. panicea, H. perlevis*), limpets (i.e., *C. fornicata*), and small crabs (i.e., *R. harrisii*), and remineralized to ^13^C-DIC. As the ^13^C-DIC concentration decreased significantly over time (Fig. 2), this bacterio-plankton-derived DIC was likely taken up by oysters (i.e., *M. gigas*) and used for shell formation. *M. gigas* is known to be able to reduce the DIC concentration in surrounding seawater during its growth period (Chen et al., 2025). In fact, the DIC concentration decreases because the formation of skeletal calcium carbonate removes the carbonate ions 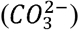 which is one of the inorganic carbon species of DIC (Liberti et al., 2022). Nitrogen from the bacterioplankton was incorporated into sponge tissue, but also re-mineralized and excreted by most organisms as ammonium. The latter was partly oxidized to nitrite by the oyster holobiont. In several incubations, the oyster holobiont likely even denitrified nitrate to nitrite as indicated by the (insignificant) negative flux of nitrate and the positive flux of nitrite (Fig. 2). This outcome agrees with studies that showed the oyster holobiont can be a hotspot of local nitrogen cycling (e.g., (Caffrey et al., 2016; Pagani et al., 2025)). For instance, in batch experiments with *M. gigas* and *Crassostrea virginica* (Caffrey et al., 2016) measured nitrification rates of 69 ± 17 nmol N cm^-2^ shell area d^-1^ for living *M. gigas* and 91 ± 20 nmol N cm^-2^ shell area d^-1^ for empty shells. In comparison, the denitrification rates of *C. virginica* were three times higher in living oysters (269 ± 37 nmol N cm^-2^ shell area d^-1^) compared to empty shells (74 ± 17 nmol N cm^-2^ shell area d^-1^) (Caffrey et al., 2016). The small negative flux of DSi (Fig. 2) indicates that the sponges used energy they gained from consuming the labile bacterioplankton to produce new sponge spicules. Spicule production requires DSi and (Reincke and Barthel, 1997) found out that the maximum silicate uptake rate of *H. panicea* is reached at a DSi concentration of 46.4 μmol l^-1^. This concentration is almost four times higher than the DSi concentration in seawater at the begin of the experiments (12.2 ± 5.17 μmol l^-1^). Hence, it is possible that the sponges consumed less DSi than expected because they were DSi-limited. Crabs enriched in ^13^C and ^15^N were not primary consumers of bacterioplankton, but took it up as secondary consumers via predation upon sponges (likely *C. maenas*) or consumption of sponge detritus (likely *E. sinensis*).

In **autumn 2020**, bacterioplankton was mostly consumed by sponges (i.e., *H. panicea*), limpets (i.e., *C. fornicata*), and small crabs (i.e., *R. harrisii*) and remineralized to ^13^C-DIC and ammonia. The latter was oxidized to nitrite and mostly nitrate, as indicated by significant releases of nitrite and nitrate (Fig. 2). As the oyster seemed to calcify less in autumn than in summer 2020 based on the DIC fluxes and therefore also grew less, it is likely that most of the nitrification in autumn happened inside the biofilm on the *M. gigas* shells. In fact, batch experiments with *C. virginica* showed that the shell biofilm excreted significant amounts of ammonia (1.26 ±0.20 μmol N ind. oyster h^-1^) and nitrite (0.05 ±0.01 μmol N ind. oyster h^-1^) (Ray et al., 2019). The shell biofilm also released most nitrate (2.17 ±1.55 μmol N ind. oyster h^-1^) when compared to the whole oyster and to the digestive system of *C. virginica*, though this flux was not significant (Ray et al., 2019). The small negative flux of DSi (Fig. 2) suggests that *H. panicea* actively took up DSi and potentially produced new spicules, despite the low ambient DSi concentration at the begin of the experiments (12.1 ± 3.59 μmol l^-1^). This spicule production could have been triggered by the food pulse in the form of ^13^C-derived bacterioplankton that was provided to the potentially food-limited *H. panicea* in autumn.

### Metabolic activity of macrozoobenthos

Following the approach of (Stratmann et al., 2025b), the metabolic activity of individual macrozoobenthos was studied via incubations with deuterium oxide. In these experiments, the specimens were considered to be most metabolically active that incorporated most ^2^H into their tissue. Hence, in summer 2020, two crab species (i.e. *C. maenas, E. sinensis*) and a sponge (i.e., *H. perlevis*) were the metabolically most active species, whereas a sponge (i.e., *H. perlevis*), a crab (i.e., *R. harrisii*), and a snail (i.e., *L. littorea*) were most metabolic active species in autumn 2020.

The metabolically very active crabs *C. maenas* and *E. sinensis* and the sponge *H. perlevis* belonged to the species that incorporated most bacterioplankton-derived ^13^C and/or ^15^N in **summer 2020**. Consequently, the metabolic activity of macrozoobenthos was directly linked to feeding activity in summer, which suggests that during this season these species might be C- and/or N-limited in the Eastern Scheldt. In fact, the primary production in the Eastern Scheldt decreased strongly between 1995 and 2009 (Smaal et al., 2013) which was interpreted as a sign for overgrazing by bivalves (Smaal et al., 2013). In addition, in summer 2019, a year before this study was conducted, *H. panicea* from the Eastern Scheldt had a low condition index which suggests that they may be starving; however, the dissolved silicate uptake data were inconclusive (Stratmann et al., 2025b). In comparison, in **autumn 2020**, the most metabolically active species were often not the species that had incorporated most bacterioplankton-derived ^13^C and/or ^15^N, except for the crab *R. harrisii*. Hence, the metabolic activity of macrozoobenthos was less linked to feeding activity than in summer. This suggests that more food was available in autumn, so that the macrozoobenthos was less food limited. Another explanation is that more species were more metabolically active in autumn that do not consume bacterioplankton. For instance, *L. littorea* is a generalist herbivore that feeds macroalgae preferably (Imrie et al., 1990). It likely grazed upon leaves of *Ulva* sp. that grew attached to the oyster shells and were more present in the autumn incubations than in the summer ones. Hence, *L. littorea* was not food limited and rather metabolically active and feeding.

A more quantitative approach to assess whether metabolic activity is linked to feeding activity may be a comparison of the 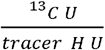 – and 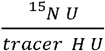 -ratios for the same species across different seasons. A larger ratio signifies that more ^13^C or ^15^N was taken up by a species that was less metabolically active (i.e., less ^2^H uptake), so the species was likely food limited, while a smaller ratio indicates that the species was metabolically more active, but took up less food, and was likely less food limited. Hence, *H. panicea*, the species with the highest 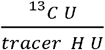 -ratio in this study, was likely more food limited in summer 2020 (median 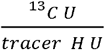 -ratio: 35.2) than in autumn 2020 (median 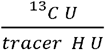 -ratio: 21.0; Fig. 4). *M. gigas*, in comparison, that is usually not the main consumer of bacterioplankton, but prefers larger particle sizes (Marks et al., 2024), was more metabolically active in summer (median 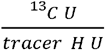 -ratio: 1.89) than in autumn (median 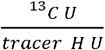-ratio: 0.39).

## Conclusion

In this study, we assessed short-term uptake of ^13^C and ^15^N-enriched bacterioplankton by *M. gigas* and its epibiota in summer and autumn 2020. Most of the bacterioplankton was taken up by filter-/suspension feeding sponges (*H. panicea* and *H. perlevis*), limpets (*C. fornicata*), and small crabs (i.e., *R. harrisii*). In contrast, other crabs took up bacterioplankton-derived ^13^C and ^15^N indirectly by predating upon *H. panicea* (*C. maenas, E. sinensis*) or taking up sponge-derived detritus (*E. sinensis*).

Furthermore, we assessed differences in the metabolic activity of *M. gigas, H. panicea*, and other associated fauna. *M. gigas* and *H. panicea* were not among the most metabolically active macrozoobenthos species, though they were more metabolically active in autumn than in summer. The most metabolically active species were crabs (*C. maenas, E. sinensis*) predating upon *H. panicea* or filter-/suspension feeding (*R. harrisii*), a sponge (*H. perlevis*), and an herbivorous snail (*L. littorea*). Hence, metabolic activity was often linked with feeding activity.

## Acknowledgments

We acknowledge the assistance of Pieter van Rijkswijk, Jeroen van Dalen, Anton Tramper (all NIOZ-EDS), and Sara Stratmann while setting-up and conducting the experiments. We thank Jan Pene and Peter van Breugel (all NIOZ-EDS), Jort Ossebar, and Ronald van Bommel (both NIOZ-MMB) for their technical support during sample processing.

This research was funded by the JPI Oceans – Ecological Aspects of Deep-Sea Mining projects under NWO-ALW grant 856.18.003 and under NWO grant NWA.1745.23.1. TS was further supported by the Dutch Research Council NWO (NWO-Rubicon grant no. 019.182 EN.012, NWO-Talent program Veni grant no. VI.Veni.212.211, NWO Open Competition Domain Science – XS grant no. OCENW.XS24.2.193) and by the European Research Council (ERC-StG grant no. 101221692, project SPYCLING).

